# Efficacy and sex differences in the effects on rat brain microglia of the colony-stimulating factor 1 receptor inhibitor-PLX5622

**DOI:** 10.1101/2022.02.22.481449

**Authors:** Aviv Sharon, Hadas Erez, Micha E. Spira

## Abstract

Microglia play pivotal roles in central nervous system development, homeostasis, responses to trauma, neurodegenerative and neuropsychiatric disorders with significant sex-bias in their symptoms and prevalence. The discovery that the survival of microglia in adult brains depends on the expression of the colony-stimulating factor 1 receptor (CSF1R), along with the development of the effective brain permeant CSF1R inhibitors PLX5622, has boosted the investigation of the role of microglia in health, disease and in relations to sex-bias. The effectiveness of PLX5622 in examining the role of microglia has mainly been demonstrated in mice. Surprisingly, despite the critical importance of rat models in brain research, there are only 4 publications in which PLX5622 was used to investigate the roles of microglia in adult rats. This has been attributed to the “impression” that PLX5622 is “ineffective in rats”. In view of the importance and interest in the role of microglia, the indispensability of rats for *in vivo* electrophysiological brain studies and behavioral research and the high efficacy of PLX5622-chow in eliminating microglia from adult mice brains, we examined the effects of PLX5622-chow on the elimination of the microglia in adult female and male rats. We found significant differences in microglia elimination by *ad libitum* PLX5622 feeding in male and female rats in different brain regions with significantly greater effectivity in female brains. Our pragmatic study provides practical information on the use and design of PLX5622 in gender-related and rat brain preclinical microglia research.

## 1. Introduction

Microglia, the resident immune cells of adult mammalian brains, play pivotal roles in central nervous system development, homeostasis, brain responses to infections or sterile insults, neurodegenerative and neuropsychiatric disorders with significant sex-bias in their symptoms, onset and prevalence (Bordt, Ceasrine, & Bilbo, 2020; Cowan & Petri, 2018; Hanamsagar et al., 2017; Kwon & Koh, 2020; Schwarz & Bilbo, 2012; Thion & Garel, 2017; Villa, Della Torre, & Maggi, 2019; Wolf, Boddeke, & Kettenmann, 2017). Under steady-state conditions, microglia morphology in adult brains is characterized by branching filopodia that dynamically “surveil” the microenvironment. Although this morphology and function is common to all “resting microglia”, specific cell geometries are seen in various brain regions (Verdonk et al., 2016). Mice microglia are characterized by slow total turnover rates of 41 months in the cortex, 15 months in the hippocampus and 8 months in the olfactory bulb (Fuger et al., 2017; Tay et al., 2017; Zhan et al., 2019), with the microglia population density maintained by local proliferation (Zhan et al., 2019). The steady-state dynamics, morphology and transcriptional phenotypes of the microglia are rapidly altered in response to different types of brain pathologies, trauma and infections (Hammond et al., 2019).

The discovery that the survival of microglia in adult brains depends on the expression of the colony-stimulating factor 1 receptor (CSF1R), along with the development of toxins, pharmacology, and gene-based approaches to blocking or eliminating the expression of CSF1R, have all enabled exploration of the role of microglia in health and disease (Green, Crapser, & Hohs, 2020; Hume et al., 2020). The recent development of effective blood-brain-barrier (BBB) permeant CSF1R inhibitors such as PLX3397 and PLX647 has advanced this field (Elmore et al., 2014; Elmore, Lee, West, & Green, 2015; Green et al., 2020). Further momentum has derived from the development of PLX5622, which exhibits higher specificity to CSF1R and improved BBB penetrance (20%) than earlier generations of brain permeant CSF1R inhibitors. Delivery of PLX5622 in rodent chow (1200mg/Kg) depleted the microglia in mouse cortex, hippocampus and thalamus by 80-90% within 5-7 days (Spangenberg et al., 2019). Use of PLX3397 in mouse chow confirmed the microglial loss as genuine and not representing downregulation of microglial markers. The microglia population recovers on withdrawal of the PLX3397 diet (Elmore et al., 2014; Elmore et al., 2015). The depletion was due to cell death and was dose- and time-dependent (Dagher et al., 2015; Elmore et al., 2014; Y. Liu et al., 2019; Najafi et al., 2018; Spangenberg et al., 2019). Immunohistological and western blot analyses revealed that the elimination of microglia by CFSR1 inhibitors is not associated with changes in the number of oligodendrocytes, neurons and astrocytes and that mice depleted of microglia show no behavioral or cognitive abnormalities (Elmore et al., 2014; Elmore et al., 2015

PLX5622 chow has been extensively and effectively used to examine the role of microglia in brain functions, maintenance, trauma and neurodegenerative diseases, mainly in mice and nonhuman-primates (over 150 studies have been published since 2015). In spite of the critical use of rats in various aspects of brain research (Ellenbroek & Youn, 2016; Homberg, Wohr, & Alenina, 2017), to the best of our knowledge there are only 4 publications on the use of PLX5622 to examine the roles of microglia in adult rats. This rather surprising fact is attributed to the “impression” that PLX5622 is “ineffective in rats” (Sehgal, Irvine, & Hume, 2021). Examination of these 4 studies raised questions as to possible sex differences in the effectiveness of PLX5622 on microglia in adult female and male rats.

Briefly, the fundamental study by Spangenberg et al. (2019; see Supplementary table 5) reported pharmacokinetic differences among mice and rats and among female and male rats in response to a single intravenous injection or one oral gavage session. This pharmacokinetic study, nonetheless, did not examine the outcome of treating adult male and female rats by PLX5622 for a number of days on microglia densities. In contrast to plausible pharmacokinetic predictions based on Spangenberg’s (2019) study, Riquier and Sollars (2020) reported that twice a day intraperitoneal injection of PLX5622 (50mg/kg) to both adult male and female Sprague Dawley rats depleted the microglia density from the gustatory system by 80–90% within 3 days, by 93% after 7 and to > 96% after 14 days of IP injections. Importantly, no sex differences were reported. In an apparent inconsistency with these findings (X. Liu, Liu, Yang, Zhou, & Tang, 2021; Riquier & Sollars, 2020) reported that in male Sprague Dawley rats 5 days of *ad libitum* PLX5622-chow (1200mg/kg) led to a small ~ 40% reduction in Iba-1 immunoblots of microglia in the spinal cord (X. Liu et al., 2021, Fig. 6), while parallel experiments on mice by the same authors showed that all the microglia were eliminated (X. Liu et al., 2021). Adding to the apparent contradictions are observations from our laboratory (Sharon, Jankowski, Shmoel, Erez, & Spira, 2021) that feeding female Sprague Dawley rats *ad libitum* with PLX5622-chow (1200mg/kg) led within 5 days to 79% elimination of cortical microglia and then further to ~94% and ~95% elimination between days 12 and 21. Taken together, the fragmented information described above cannot serve as a solid background for designing studies in which PLX5622 is used in rats to decipher basic and preclinical aspects of microglia function.

There is growing interest in the roles of microglia in normal and pathological brains and in the gender-bias in the prevalence, severity and onset of neuro-disorders related to microglia. The high efficacy of PLX5622-chow (1200mg/kg) in eliminating microglia from adult mice brains and the extensive use of rats as a model system for behavioral and *in vivo* electrophysiological brain research led us to examine the effects of PLX5622 chow (1200 PPM) on the elimination of the brain microglia population in female and male rats. Our pragmatic study provides practical information on the use and design of PLX5622 in gender-related and rat brain preclinical microglia research.

## 2. Materials and Methods

### 2.1 Animals

12 week old male and female Sprague Dawley rats from the same litter were used for all experiments. All procedures were approved by the Committee for Animal Experimentation at the Institute of Life Sciences of the Hebrew University of Jerusalem. All procedures were carried out in accordance with the approved guidelines. The immunohistological studies were conducted using female and male Sprague Dawley rats (240-340 g). For microglial ablation, the rats were fed ad libitum, for 10 days, with a PLX5622 diet (1200 PPM PLX5622, Plexxikon Inc., Berkeley, USA). PLX5622 was provided by Plexxikon Inc. and formulated in AIN-76A standard chow by Research Diets Inc.

### 2.2 Tissue Processing for Immunohistology

Control rats and rats fed *ad libitum* with PLX5622-chow for ten days were sacrificed for immunohistological examinations of the microglia. To that end, Individual rats were deeply anesthetized with isoflurane (Piramal, United States) followed by an IP overdose injection of Pentel (4.5 ml per 250 g rat, CTS Group, Israel). When the rats stopped breathing, they were transcardially perfused with phosphate buffer saline (PBS) at a rate of 10 ml/min for 40 min. This was followed by perfusion with 4% paraformaldehyde in PBS (PFA, Sigma-Aldrich) at a rate of 10 ml/min for 40 min. Next, the skulls was removed and the brain was post-fixed at 4°C for an additional 12–24 h. in 4% PFA. Afterwards, brains were washed in PBS and incubated for 1–3 days in a 30% sucrose solution in PBS at 4°C.

### 2.3 Cryosectioning and Immunohistological Labeling

To prepare for cryosectioning, cubes of tissue were isolated from different brain regions. These were placed in a freezing medium (Tissue-Plus O.C.T. Compound, Scigen) and frozen at −80°C. The frozen tissue was then sectioned at 40 μm in the coronal plane using a Leica CM1850 Cryostat. Individual slices were collected and placed in 24 well plates containing PBS.

The tissue slices were then incubated in blocking solution (1×PBS, 1% heat-inactivated horse serum (Biological Industries), 0.1% Triton X-100 (Sigma Aldrich)) for 1 hour at room temperature (RT) under gentle shaking. Next, the slices were incubated with a diluted primary antibody for 3 hours at RT and washed 3 times with the blocking solution. This was followed by a 1 hour incubation at RT with the diluted secondary antibody, after which the slices were washed with the blocking solution 3 times and stained with the nuclear marker DAPI (Sigma–Aldrich, 1 mg/ml 1:1000) for 15 min at RT. After washing with the blocking solution and PBS the slices were mounted on Superfrost Plus Slides (Thermo Fisher Scientific) and sealed with a Vectashield (VE-H-1000 - Vector Labs) mounting medium. Meticulous examination of the prepared tissue slices by confocal microscope optical sectioning revealed that the antibodies penetrated the tissue to homogeneously stain the target cells (Huang et al., 2020; Sharon et al., 2021).

Microglia were labeled using rabbit anti-Ibl-1 monoclonal antibody (Abcam ab178846, 1:2000). For the secondary antibodies we used sheep anti-rabbit Cy3 (Sigma–Aldrich C2306, 1:100). To confirm that the Iba-1 antibody labeled resident microglia and not infiltrated macrophages we co-labeled the Iba-1 positive cells using goat anti Iba-1 antibody (Abcam ab5076, 1:125) with rabbit polyclonal antibody for rat TMEM119 that recognizes microglia-specific transmembrane proteins (Synaptic Systems GmbH, 1:1000, 400 203). For the secondary antibodies, we used donkey anti-goat Igg H&L (Alexa Fluor® 647) Preadsorbed (Ab150135, 1:50) and sheep anti-rabbit Cy3 (Sigma–Aldrich C2306, 1:100).

### 2.4 Microscopy

Confocal image stacks of the immunolabeled slices were acquired with an Olympus FLUOVIEW FV3000 confocal scan head coupled to an IX83 inverted microscope, using a 20× air objective (NA = 0.75). Sections were scanned in sequential mode. DAPI (not shown) was excited using the 405nm laser and its emission was acquired in the 415-470nm range using a spectral detector, Cy3 was excited using the 561nm laser and its emission was acquired in the 570–630nm range using a second spectral detector. A non-confocal transmitted light image was also acquired.

For the co-staining of Iba1 and TMEM-119 (not shown) we used sequential mode: Alexa 647 was excited using the 640nm laser and its emission was acquired in the 645-745nm range using a spectral detector, Cy3 was excited using the 561nm laser and its emission was acquired in the 575–650nm range using a second spectral detector. A non-confocal transmitted light image was also acquired.

Typically, 15-35 confocal slices were acquired with a vertical spacing of 1μm. Image stacks were acquired from 6 different brain areas: cortex, hippocampus, amygdala, striatum, cerebellum and olfactory bulb. For microglia morphology analysis we set a x2 zoom.

### 2.5 Image Processing, Analysis and Statistics

The Image processing was implemented using the Fiji distribution of ImageJ (Schindelin et al., 2012; Schneider, Rasband, & Eliceiri, 2012), as follows. A maximum intensity projection image was created using 10 consecutive optical sections from each of the 40μm thick brain slices. The cells were counted manually from each image. The sample size of the immunohistological sections is given in Supplementary Table 1. Significant statistical differences between the different treatments (females and males, plx5622 and control diet) were determined by a t-test for two samples assuming unequal variances. For all tests, a P value < 0.01 indicated a statistically significant difference (Supplementary Table 2).

### 2.6 Microglia’s morphology analysis

A maximum intensity projection image was created using the Fiji distribution of ImageJ (Schindelin et al., 2012; Schneider et al., 2012). All consecutive optical sections from each of the 40-μm-thick brain slices were used (from stacks of x 2 zoom). The file was saved as an 8 bit gray image (e.g., see Supplementary Figure 1A).

Those images were imported to the HCA-vision software (CSIRO Australia, Vallotton et al., 2007; Wang et al., 2010). The microglia cell bodies and extensions were automatically identified and measured (Supplementary Figure S1). The parameters chosen for the analysis of the microglia morphology were: (a) Number of roots - the number of points where an extension structure touched a microglia cell body. (b) Number of branch points - the number of points where a microglia extension structure split into 2 or more branches. (c) Number of segments - a segment is a linear structure between branching points or a microglia cell body. (d) Number of extremities - the number of terminating extension segments. (e) Total extension length - sum of the length of all extensions.

For the analysis we chose only cells that were properly recognized by the software and whose extensions were not cut at the edges of the image. For the automated microglia morphology analysis we used the same optimal parameters setting for each group (male, female, control and PLX5622 fed rats). The quality of the automated detection by these parameters setting is presented in Supplementary Figure S1. The final data and group sample size are given in Supplementary Table 1.

## 3. Results

### 3.1 Steady-state densities and morphological phenotypes of microglia in the brains of female and male rat

A number of studies demonstrated that in adult mice the microglia densities and morphology differ in different brain regions and among females and males (Guneykaya et al., 2018; Lenz, Nugent, Haliyur, & McCarthy, 2013; McCarthy, Pickett, VanRyzin, & Kight, 2015; Mouton et al., 2002; Schwarz & Bilbo, 2012; Tan, Yuan, & Tian, 2020).

To gain missing analogues information in rat brains, we mapped microglia densities and morphological phenotypes in adult female and male Sprague Dawley rats using Iba-1 antibody immunohistological labeling (Figure 1). Because of economic considerations we conducted the analysis on rats fed with PLX5622 chow for 10 days. This point in time was selected as in an earlier study (Sharon et al 2021), we found that maximal microglia elimination is achieved in female rats between 7 to 12 days of PLX5622 feeding. The immunohistological protocol was confirmed by co-labeling Iba-1 positive cells with rat TMEM119 antibody that recognizes microglia-specific transmembrane proteins (e.g., Huang et al., 2020; Sharon et al., 2021). Microglia densities in the cortex, hippocampus, amygdala, striatum, cerebellum and olfactory bulb were mapped using stacks of confocal microscope images and the HCA-Vision/Acapella software.

**Figure 1.**
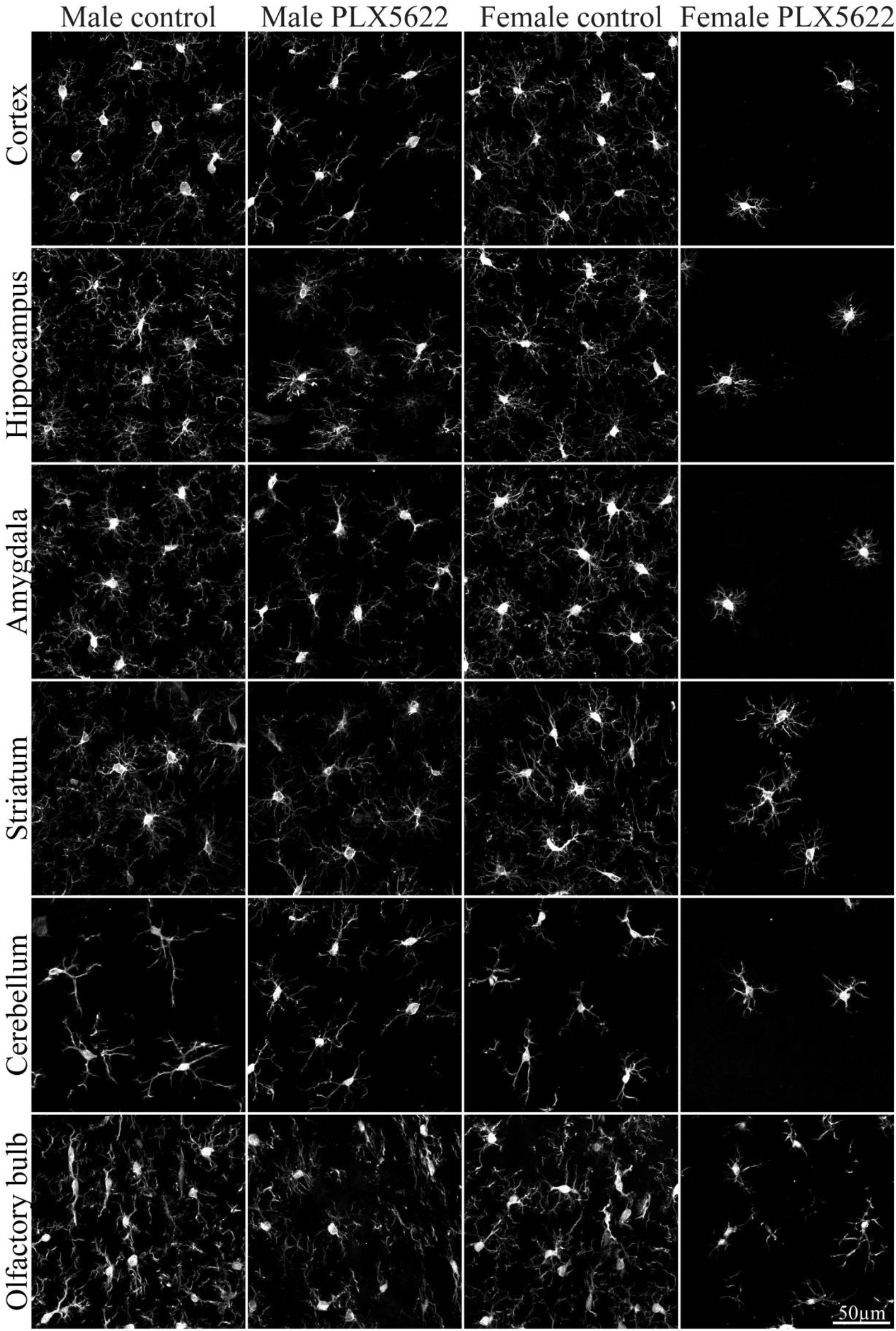
Confocal microscope images of Iba-1 labeled microglia in different brain regions in control male and female rats and rats fed *ad libitum* for 10 days on PLX5622 chow.

As reported for mice brains (Tan et al., 2020) differences in microglia densities were seen in different regions in the rat brain (Figure 2, Supplementary Tables 1, 2). In addition, some of these regions showed small but significant sex differences (Figures 2, Supplementary Table 2, Guneykaya et al., 2018; Wolf et al., 2017). As in mice, in both female and male rats, the lowest microglia densities were found in the cerebellum (235±17cells/mm^2^ in males and 228±25 mm^2^ in females) and the highest in the olfactory bulb (391±36 cells/ mm^2^ in males and 381±34 mm^2^ in females).

**Figure 2.**
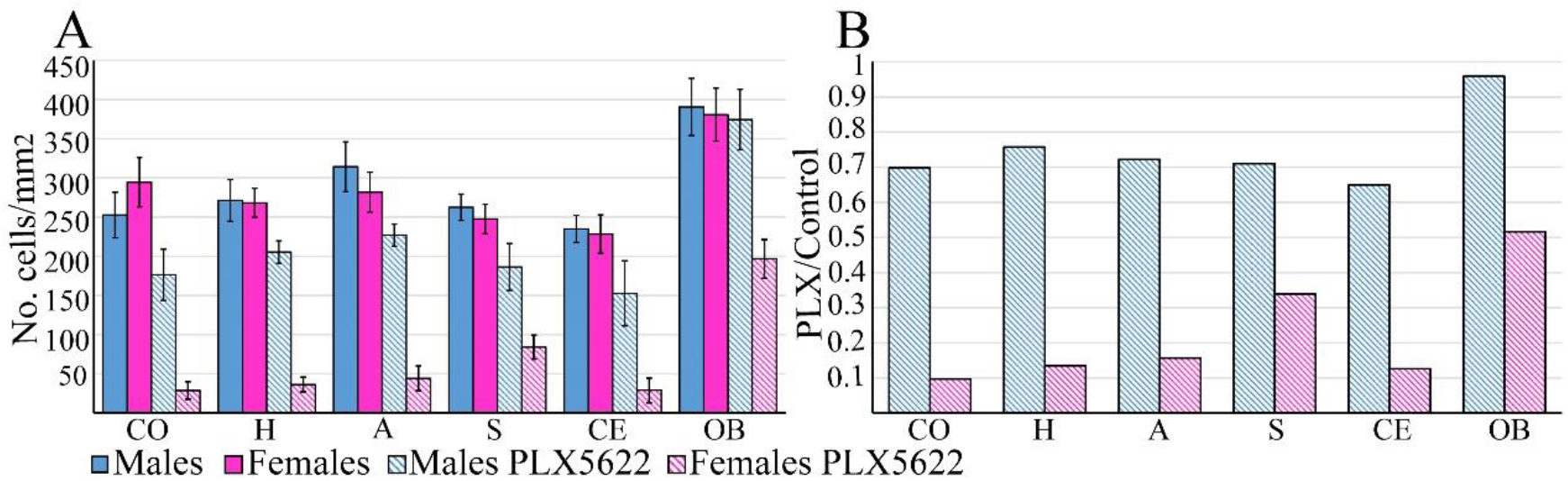
**(A) The average microglia densities (number of cells/mm**^**2**^**) in different brains regions in control (homogeneous columns) male (blue) and female (magenta) rats, and rats fed for ten days *ad libitum* on PLX5622 chow (diagonal stripes)**. Controls showed small but significant density differences between females and males within given brain regions. Large and significant differences in microglia densities were seen between females and males in different brain regions among rats fed on PLX5622. In both males and females, the lowest density of microglia was in the cerebellum, the highest in the olfactory bulb. Large and significant differences in the elimination of microglia from males and females were observed in all brain regions in response to PLX5622 chow (Supplementary Table 2). Note that the in male olfactory bulb the density of microglia was not altered by PLX5622. (B) Ratio values of averaged microglia densities in PLX5622-treated rats and control rats in different brain regions showed a significantly smaller effect of PLX5622 chow on male microglia densities than in females. CO-cortex, H-hippocampus, A-amygdala, S-striatum, CE-cerebellum, OB-olfactory bulb.

Earlier studies documented that, at steady state, microglia in all brain regions extend delicate ramifying branches (Figures 1, 3). Yet, quantitative morphological analysis of mouse microglia revealed characteristic morphological differences in different brain regions (De Biase & Bonci, 2019; Fernandez-Arjona, Grondona, Granados-Duran, Fernandez-Llebrez, & Lopez-Avalos, 2017; Tan et al., 2020).

**Figure 3.**
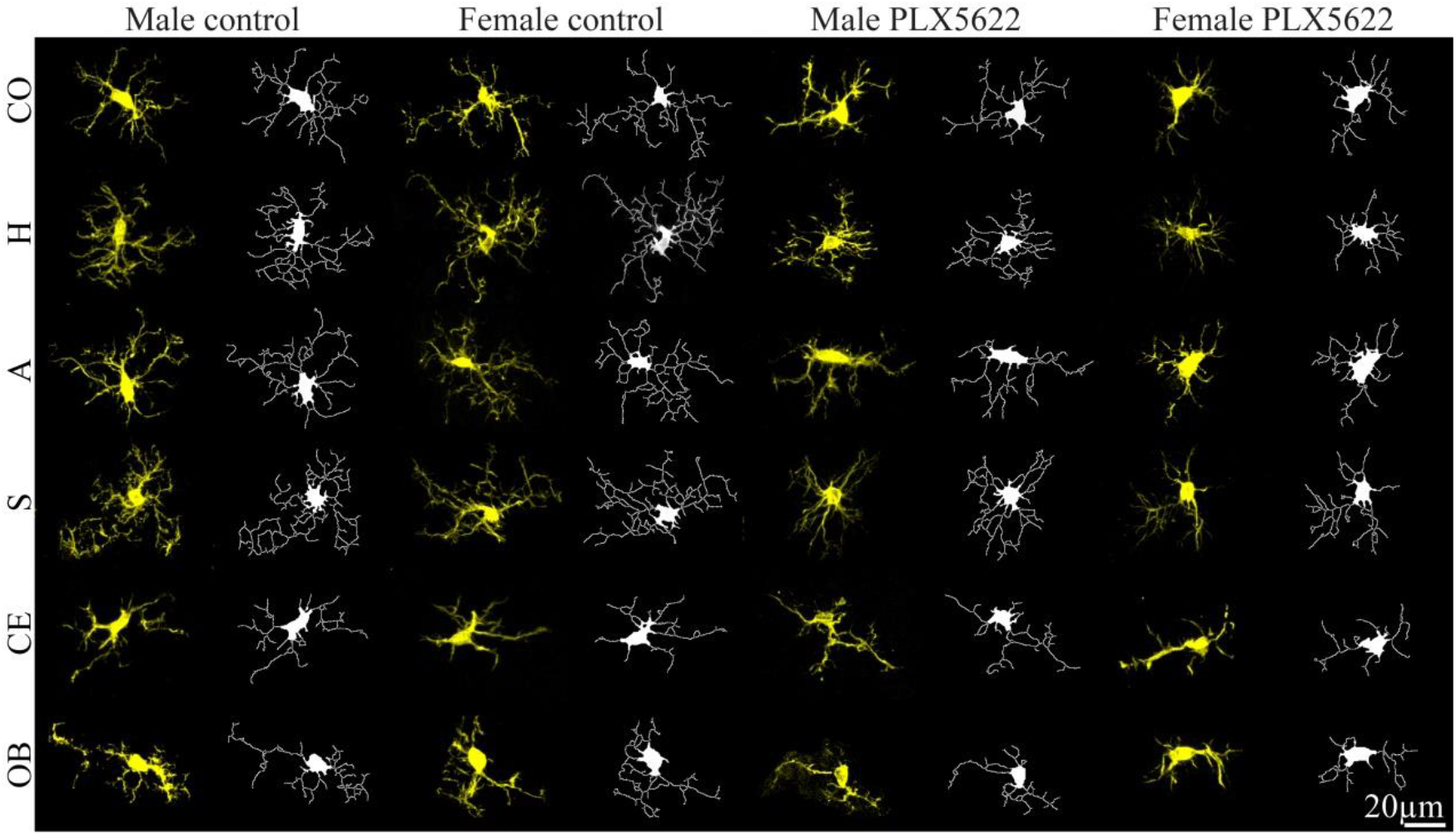
Maximum intensity projection confocal microscope images (yellow) from different brain regions in control male and female and in rats fed *ad libitum* for 10 days on PLX5622 chow. These were imported to the HCA-vision software (white) and the microglia cell bodies and extensions were automatically identified and measured. CO-cortex, H-hippocampus, A-amygdala, S-striatum, CE-cerebellum, OB-olfactory bulb (for statistics see Supplementary tables 1,2).

Our analysis of microglia shown in Figures 3,4 confirmed morphological differences in different brain regions and among male and female rats. The average number of branches extending from a single cell body (roots/cell) was similar in males and females in all examined regions but was significantly smaller in the cerebellum and olfactory bulb (Supplementary Table 2). The averaged total microglia branch length, the number of segments, branching points and extremities was similar among males and females in the cortices, striatum and the olfactory bulbs (Figure 4, Supplementary Table 2) but there were sex differences in other brain regions.

**Figure 4.**
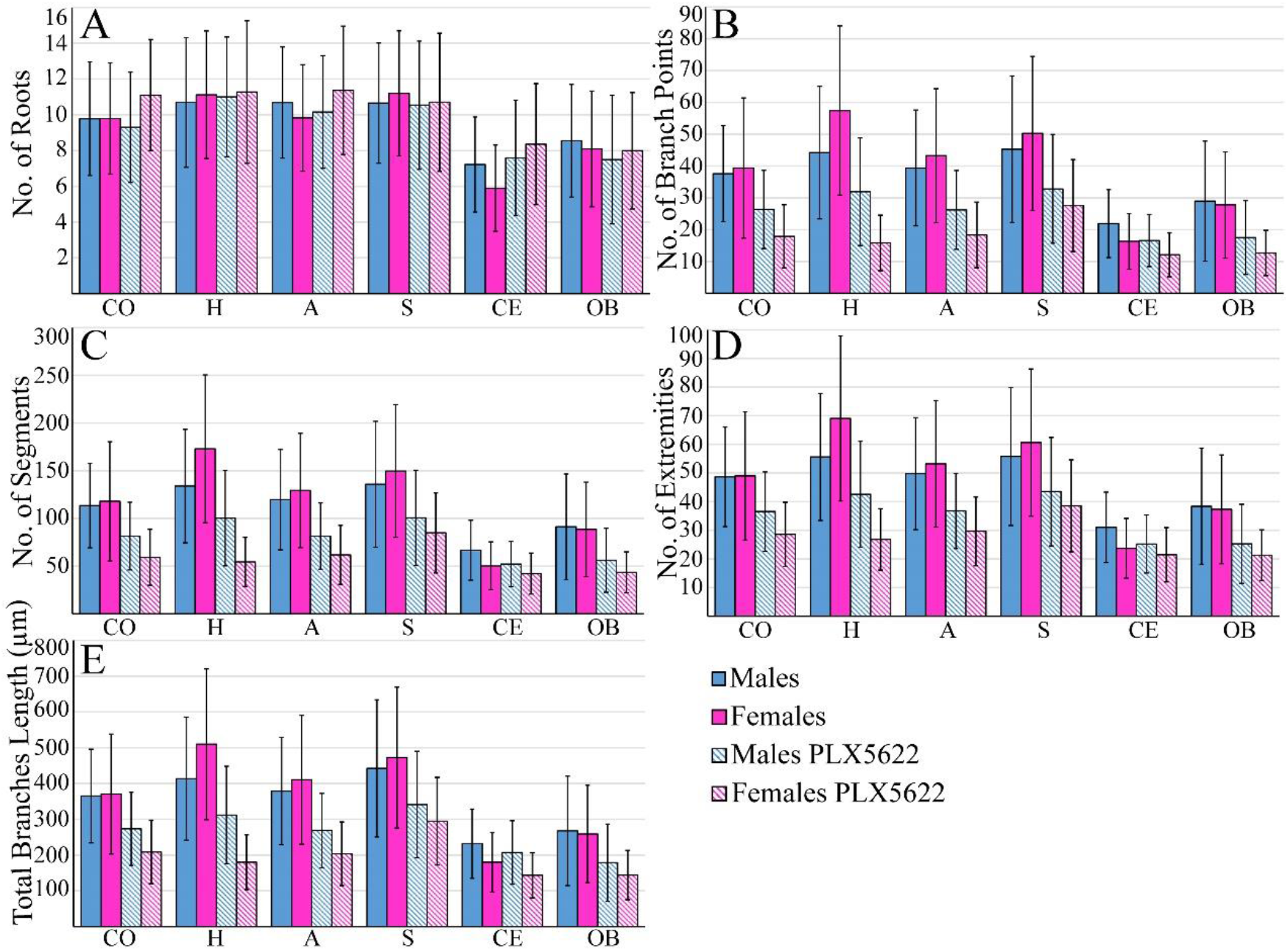
Quantitative morphological analysis of microglia in different brain regions in control (homogeneous columns) male (blue) and female (magenta) rats and rats fed *ad libitum* for ten days on PLX5622 chow (diagonal stripes). (A) The number of roots/cell, (B) the number of branching points, (C) number of segments, (D) number of extremities and (E) total branch lengths. The number of roots/microglia in control and microglia surviving PLX5622 in the cerebellum and olfactory bulb was lower than in other brain regions and was similar in males and females in all brain regions examined and was unaltered by PLX5622 treatment. Microglia surviving the 10 days of PLX5622 chow showed a reduced number of branching points, segments, extremities and total branch length in all examined brain regions. CO-cortex, H-hippocampus, A-amygdala, S-striatum, CE-cerebellum, OB-olfactory bulb.

### 3.2 PLX5622 induced microglia elimination and morphological alterations of the surviving microglia

The effects of PLX5622 chow on microglia density in mice showed that microglia elimination is dose-dependent. For example *ad libitum* feeding mice with 300 mg PLX5622/kg chow resulted in sustained 30% elimination of the microglia whereas 1200 mg/kg PLX5622 chow resulted in 80% elimination within a week. The conclusion was that “elimination is not an all or nothing event but something that can be modulated” and that males and females mice respond equally to the drug (Dagher et al., 2015).

Our study showed that the microglia densities in both female and male rats were generally reduced following 10 days of *ad libitum* feeding with PLX5622 chow (1200mg/kg). In **<u>female</u>** cortices, hippocampi, amygdala and cerebellum the microglia density was significantly reduced to 0.1-0.16 of control values (Figure 2, Supplementary Tables 1, 2), to 0.34 in the striatum, and to 0.52 in the olfactory bulb (Figures 1 and 2). In **<u>males</u>**, the microglia density was significantly less reduced only to 0.65-0.76 of the control value in the cortices, hippocampi, amygdala, striatum and cerebellum and was unchanged in the olfactory bulb (Figures 1, 2, Supplementary Tables 1 and 2).

Since the elimination of the microglia by CSF1R inhibitors is known to be dose-dependent, and as the effect of PLX5622 chow on males and females rats differed, we examined whether the microglia surviving after 10 days of PLX5622 feeding in both sexes had undergone any structural changes and if so, whether the changes were similar. Morphological analysis of these microglia revealed that, except for the number of roots/cell and except for the cerebellum, all other parameters (number of branches, segments, extremities and branch length) were reduced in relation to the control in both females and males (Figures 4, 5, Supplementary Tables 1, 2). The morphological changes were more pronounced in females than in males, but the averaged differences among males and females was small.

**Figure 5.**
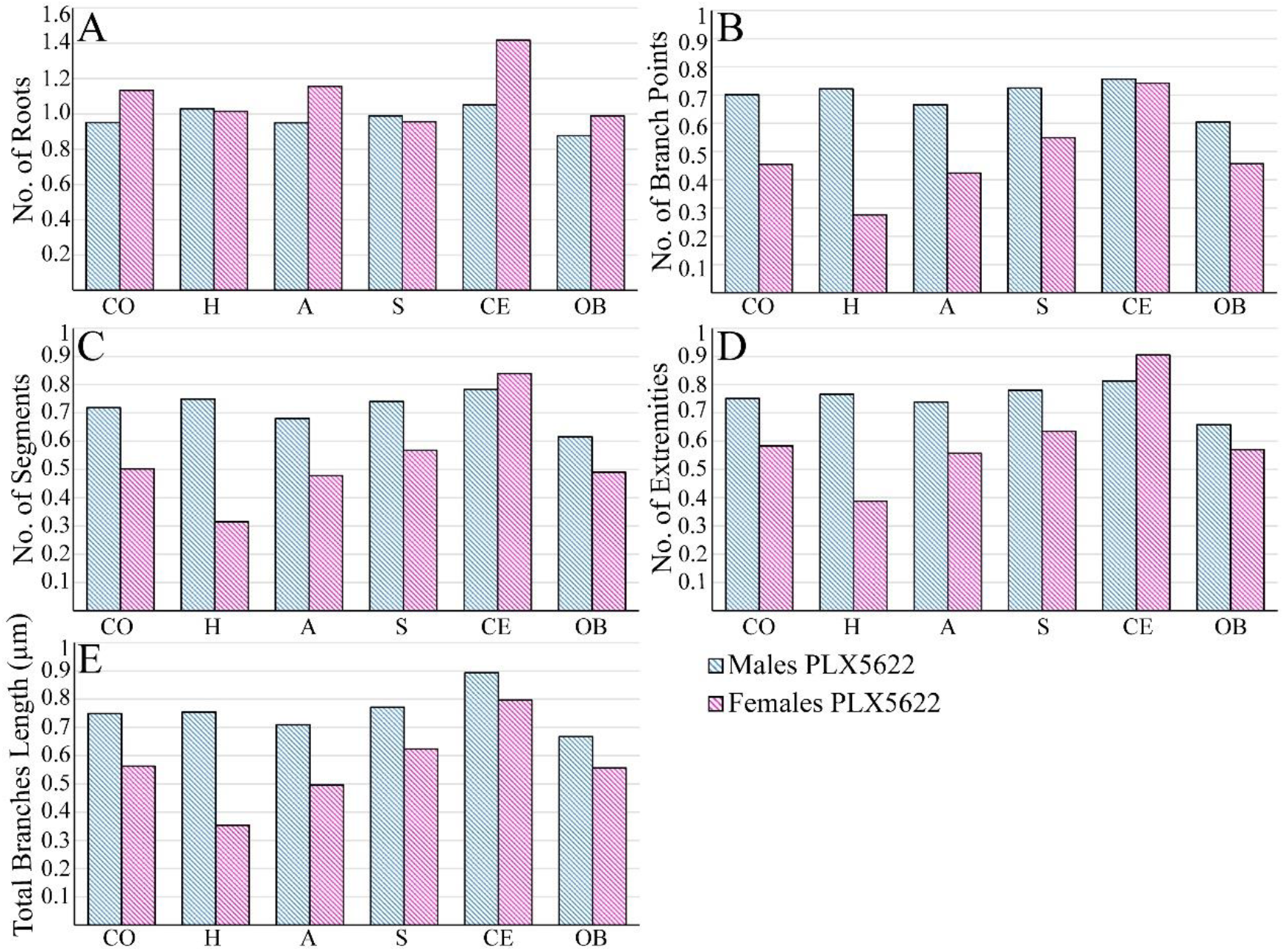
Ratio values of the numbers of (A) roots/cell, (B) branching points, (C) segments, (D) extremities and (E) total branch lengths in microglia surviving 10 days of PLX5622-chow compared to the control rats in different brain regions. In females microglia surviving PLX5622 showed a reduction in all parameters and brain regions except for the number of roots/microglia and at the cerebellum. CO-cortex, H-hippocampus, A-amygdala, S-striatum, CE-cerebellum, OB-olfactory bulb.

In conclusion, the morphological analysis of the microglia that survived, a 10 days period of *ad libitum* PLX5622 feeding revealed that beside for the number of roots/cell that were not altered (Figure 4), all other parameters (branching pattern and branches length) in both females and males were reduced in respect to controls.

## 4. Discussion

Delivering CSF1R inhibitor PLX5622 in rodent chow is advantageous for studying the roles of microglia in health and pathological conditions. PLX5622 chow has shown a dose-dependent effect in reducing microglia density in mice; “Elimination is not an all or nothing event but something that can be modulated”; male and female mice respond equally to the drug (Dagher et al., 2015). In view of the advantageous use of the CSF1R inhibitor PLX5622 in the form of rodent chow in studying the roles of microglia in health and pathological conditions, and because of uncertainties as to the effects of PLX5622 on adult rat microglia (Riquier & Sollars, 2020; Sehgal et al., 2021; Sharon et al., 2021; Spangenberg et al., 2019), we examined two questions (a) Are there any limitations in the use of adult rats as basic and preclinical models for the study of microglia using the CSF1R inhibitor PLX5622 in chow and (b), are there sex differences in the response of rat brains to PLX5622 chow treatment? These pragmatic questions are important, as rats are preferable to mice models for in vivo electrophysiological brain studies and behavioral studies, mainly because of their body size. PLX5622 chow is also a very convenient experimental tool for introducing an effective CSF1R antagonist (for example: Hume et al., 2020; Patkar et al., 2021; Schwarz & Bilbo, 2012).

Our main observations were that at steady-state, the average microglia density was similar in female and male rat brains (although the small differences observed in the cortex, amygdala and striatum were statistically significant; P≤0.01, Supplementary Table 2). The rate of PLX5622 chow-induced microglia elimination in female rats was somewhat slower than that in mice (Sharon et al., 2021). Nonetheless, microglia elimination in female rats proceeded within 10 days to 0.1-0.16 of the control levels in cortices, hippocampus, amygdala and cerebellum, in the striatum to 0.34, and only to 0.52 in the olfactory bulb. In contrast, 10 days of *ad libitum* PLX5622-chow feeding of male rats did not lead to reduction in the average microglia density in the olfactory bulb (0.96) and there was significantly less reduction than in females (0.65-0.76) in other regions of the brain (Figures 1, 2, Supplementary Tables 1,2).

The sex-related differences in the effects of PLX5622-chow on microglia densities could reflect pharmacokinetic differences among females and males, as suggested by single intravenous injection or single oral gavage of PLX5622 by Spangenberg et al. (2019). Since the microglia densities in male rat brain (apart from the olfactory bulb) were reduced by PLX5622 feeding (Figures 1, 2), and as the structural phenotypes of the surviving microglia were altered similarly to those in females (Figures 3-5) it is conceivable to assume that PLX5622 provided in the chow did cross male rat BBB to reach the brain parenchyma. Since the effect of CSF1R inhibition on microglia survival is dose-dependent (Dagher et al., 2015), the observed sex differences in the number of eliminated microglia may reflect differences in the effective PLX5622 concentration in the brain parenchyma of females and males fed *ad libitum*. Riquier and Sollars (2020) explicitly stated that no gender differences were noticed in microglia elimination in the rat gustatory system following intraperitoneal injections of PLX5622 in adult rats. It is thus reasonable to assume that the introduction of PLX5622 by intraperitoneal injection protocol rather than by feeding bypass undefined pharmacokinetic barriers to elevate the effective PLX5622 brain concentration to a level that blocked the CSF1R in both males and females. Nonetheless, since differences in microglia elimination levels in some brain regions were significantly larger than in others (e.g., in male olfactory bulb microglia were not eliminated while in other regions the population was reduced to 0.65-0.76 of control and in females, microglia elimination level in the olfactory bulb was smaller than in other regions, Figure 2), we cannot rule out the possibility that some differences in CSF1R densities, receptors affinity to PLX5622 or differences in downstream molecular cascades of CSF1R activation in male rats could account for the gender differences From a pragmatic point of view, we conclude that adult female rats and PLX5622-chow can be effectively used to examine the *in vivo* role of microglia in brain function. Further studies are needed to uncover the mechanisms underlying the sex differences.

## Acknowledgments

This work was supported by the Israel Science Foundation grant number 1808/19. Part of this work was conducted at the Charles E. Smith and Prof. Joel Elkes Laboratory for Collaborative Research in Psychobiology. This study is based on an earlier research project supported by the National Institute of Neurological Disorders and Stroke of the National Institutes of Health under Award Number U01NS099687.

## Author contributions

A.S. implanted the platforms and processed the tissues for immunohistological sectioning. A.S. and H.E. analyzed the images. M.E.S. conceived, designed, and supervised the project.

