## Supplemental data for "Efficacy and sex differences in the effects on rat brain microglia of the colony-stimulating factor 1 receptor inhibitor-PLX5622"

**Supplementary material: 22 02 2022**

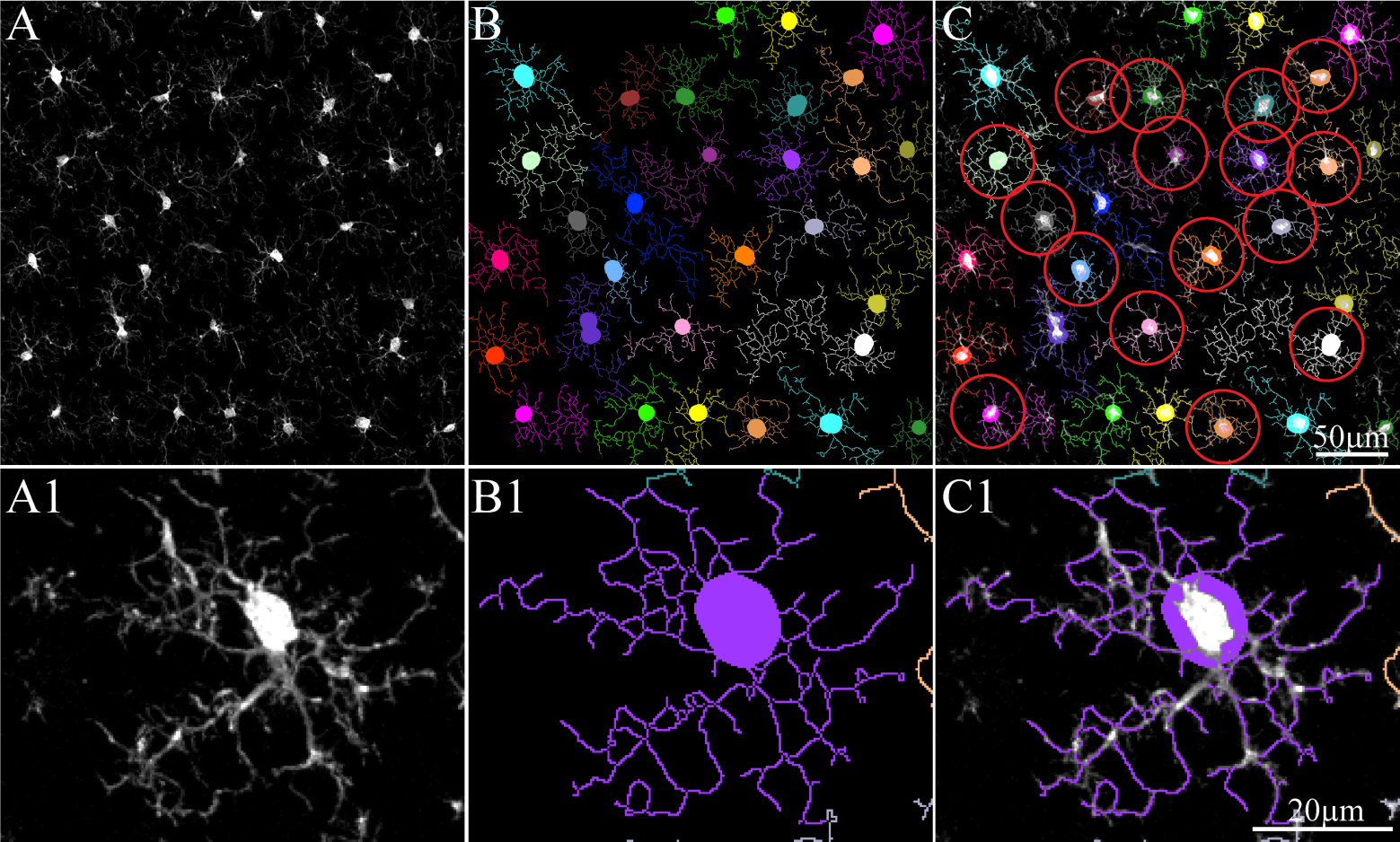

**Supplementary Figure S1.** Sequence of microglia morphological analysis. (A, A1) Maximal intensity projection images produced from z-stacks taken by confocal microscope. (B, B1) The cell bodies and extensions detected by HCA-vision software. (C, C1). Merge image of A/A1 and B/B1. Only the cells marked by red circle were taken for the final analysis. Cells that were cut at the edges of the image were not included in the analysis.

**Supplementary Table1: Density and morphology sample size, means and standard deviations of microglia in male and female rats.**

| **Brain's Area** | **Group** | **Group size (Density)*** | **Density (Number of cells/mm2)** | **Group size (Morphology)*** | **Total Neurite Length (µm)** | **Number of Branch Points (B)** | **Number Of Roots (R)** | **Number Of Segments (S)** | **Number Of Extremities**  **(E)** |
| --- | --- | --- | --- | --- | --- | --- | --- | --- | --- |
| **Cortex** | Male | 4H, 30S | 252.7±29.1 | 20S, 247C | 365±130.9 | 37.6±15.1 | 9.8±3.2 | 113.5±44.2 | 48.6±17.4 |
|  | Male PLX5622 | 4H, 20S | 176.4±32.6 | 18S, 227C | 273.3±102.2 | 26.4±12.3 | 9.3±3.1 | 81.6±35.5 | 36.5±13.9 |
|  | Female | 4H, 28S | 294.5±31.5 | 18S, 236C | 370.5±167.5 | 39.4±22 | 9.8±3.1 | 118±62.5 | 49±22.4 |
|  | Female PLX5622 | 4H, 18S | 28.7±11.2 | 96S, 217C | 208.6±88.6 | 17.9±9.9 | 11.1±3.1 | 59.3±29.5 | 28.6±11.3 |
| **Hippocampus** | Male | 4H, 20S | 271.2±26.6 | 20S, 202C | 413.4±172.2 | 44.2±20.8 | 10.7±3.6 | 134±59.4 | 55.6±22.2 |
|  | Male PLX5622 | 4H, 20S | 205.4±14.3 | 20S, 166C | 311.8±136.4 | 31.9±16.9 | 11±3.3 | 100.3±50 | 42.6±18.5 |
|  | Female | 4H, 20S | 268.3±18.4 | 20S, 230C | 509.7±210.7 | 57.4±26.6 | 11.1±3.6 | 173±77.5 | 69.1±28.8 |
|  | Female PLX5622 | 4H, 20S | 36.2±9.4 | 80S, 138C | 180.2±76.8 | 15.8±8.7 | 11.3±4 | 54.4±25.8 | 26.8±10.7 |
| **Amygdala** | Male | 4H, 20S | 314.3±31.7 | 20S, 379C | 378.7±149.8 | 39.4±18.2 | 10.7±3.1 | 119.7±52.7 | 49.8±19.6 |
|  | Male PLX5622 | 4H, 20S | 227±14.1 | 19S, 243C | 268.6±103.8 | 26.2±12.4 | 10.2±3.2 | 81.4±34.7 | 36.7±13.2 |
|  | Female | 4H, 20S | 281.9±25.4 | 19S, 324C | 410.5±180.1 | 43.3±21.1 | 9.8±3 | 129.4±59.9 | 53.2±22.1 |
|  | Female PLX5622 | 4H, 20S | 44.2±15.8 | 72S, 200C | 203.5±88.9 | 18.3±10.3 | 11.4±3.6 | 61.9±31 | 29.6±12 |
| **Striatum** | Male | 4H, 20S | 262.6±16.8 | 20S, 233C | 442.3±191.8 | 45.3±23 | 10.7±3.4 | 135.9±66 | 55.8±24.1 |
|  | Male PLX5622 | 4H, 20S | 186.4±30 | 20S, 219C | 341.1±148.9 | 32.8±17.1 | 10.5±3.6 | 100.6±49.9 | 43.5±18.9 |
|  | Female | 4H, 18S | 247.6±18.8 | 20S, 176C | 472.3±196.9 | 50.3±24.2 | 11.2±3.5 | 149.7±69.5 | 60.6±25.7 |
|  | Female PLX5622 | 4H, 19S | 84±15.4 | 80S, 421C | 294.6±122.1 | 27.6±14.4 | 10.7±3.9 | 85±42 | 38.5±16.1 |
| **Cerebellum** | Male | 4H, 19S | 235±17.2 | 20S, 181C | 231.9±96.9 | 21.9±10.7 | 7.2±2.7 | 66.7±31.4 | 31±12.3 |
|  | Male PLX5622 | 4H, 20S | 152.6±41.6 | 20S, 141C | 207.2±88.8 | 16.6±8.2 | 7.6±3.2 | 52.3±23.7 | 25.2±10.1 |
|  | Female | 4H, 16S | 228.4±25.4 | 20S, 196C | 180±82.9 | 16.3±8.7 | 5.9±2.4 | 50.3±25.1 | 23.7±10.4 |
|  | Female PLX5622 | 4H, 20S | 28.8±15.8 | 80S, 153C | 143.4±63.1 | 12.1±6.9 | 8.4±3.4 | 42.2±21.3 | 21.5±9.5 |
| **Olfactory bulb** | Male | 4H, 19S | 390.6±36.5 | 19S, 322C | 267.7±153 | 29±18.9 | 8.5±3.1 | 91.3±55.4 | 38.4±20.3 |
|  | Male PLX5622 | 4H, 20S | 374.4±38.6 | 20S, 268C | 178.6±107.7 | 17.5±11.6 | 7.5±3.6 | 56.2±33.5 | 25.3±13.8 |
|  | Female | 4H, 18S | 380.8±33.8 | 19S, 330C | 259±136.3 | 27.8±16.6 | 8.1±3.2 | 88.6±49.4 | 37.3±19 |
|  | Female PLX5622 | 4H, 19S | 196.6.1±24.8 | 46S, 353C | 144.1±68.8 | 12.7±7.1 | 8±3.3 | 43.4±21.4 | 21.3±8.9 |

***** H= hemispheres, S=slices, C= cells

**Supplementary Table 2: T tests of microglia density and morphology in male and female rats (P>0.01, P<0.01).**

| **Density** | |  | |  | |  | |  |
| --- | --- | --- | --- | --- | --- | --- | --- | --- |
| **Area** | | **Female control versa Male control** | | **Female control versa Female PLX** | | **Male control versa Male PLX** | | **Female PLX versa Male PLX** |
| **Cortex** | | 1.28232E-06 | | 4.93603E-32 | | 1.47044E-10 | | 2.7351E-16 |
| **Olfactory bulb** | | 0.199825468 | | 8.10872E-19 | | 0.09289157 | | 2.38809E-18 |
| **Cerebellum** | | 0.188266419 | | 3.15884E-20 | | 6.20764E-09 | | 2.88576E-12 |
| **Hippocampus** | | 0.343538842 | | 2.72361E-29 | | 5.68626E-11 | | 2.59566E-31 |
| **Amygdala** | | 0.000509723 | | 1.32327E-27 | | 8.67007E-12 | | 2.03966E-32 |
| **Striatum** | | 0.00727998 | | 2.66118E-25 | | 2.90375E-11 | | 2.3705E-14 |
| **Total branches Length** | |  | |  | |  | |  |
| **Area** | | **Female control versa Male control** | | **Female control versa Female PLX** | | **Male control versa Male PLX** | | **Female PLX versa Male PLX** |
| **Cortex** | | 0.342899156 | | 2.53799E-32 | | 1.03102E-16 | | 1.94794E-12 |
| **Olfactory bulb** | | 0.222209133 | | 7.67028E-37 | | 4.71782E-16 | | 3.02175E-06 |
| **Cerebellum** | | 2.56074E-08 | | 2.05095E-06 | | 0.008992885 | | 8.77326E-12 |
| **Hippocampus** | | 1.3905E-07 | | 5.72807E-64 | | 3.78793E-10 | | 2.2061E-22 |
| **Amygdala** | | 0.00607278 | | 2.98931E-54 | | 2.1957E-25 | | 2.25042E-12 |
| **Striatum** | | 0.06143642 | | 1.29616E-23 | | 4.02957E-10 | | 4.16974E-05 |
| **Number Of Roots** | |  | |  | |  | |  |
| **Area** | | **Female control versa Male control** | | **Female control versa Female PLX** | | **Male control versa Male PLX** | | **Female PLX versa Male PLX** |
| **Cortex** | | 0.48466279 | | 5.03248E-06 | | 0.047230387 | | 1.03961E-09 |
| **Olfactory bulb** | | 0.03346394 | | 0.353285939 | | 9.88681E-05 | | 0.036755492 |
| **Cerebellum** | | 1.49724E-07 | | 2.26186E-13 | | 0.13686744 | | 0.023037164 |
| **Hippocampus** | | 0.108314615 | | 0.351139994 | | 0.195996501 | | 0.259877645 |
| **Amygdala** | | 9.88841E-05 | | 3.13381E-07 | | 0.019802837 | | 0.000112395 |
| **Striatum** | | 0.058135284 | | 0.062810578 | | 0.359193931 | | 0.299095526 |
| **Number of Branch Points** | |  | |  | |  | |  |
| **Area** | | **Female control versa Male control** | | **Female control versa Female PLX** | | **Male control versa Male PLX** | | **Female PLX versa Male PLX** |
| **Cortex** | | 0.149955075 | | 6.82347E-34 | | 4.67864E-18 | | 7.02304E-15 |
| **Olfactory bulb** | | 0.191980437 | | 1.06921E-42 | | 1.3208E-18 | | 1.70569E-09 |
| **Cerebellum** | | 3.58128E-08 | | 3.77789E-07 | | 3.44706E-07 | | 4.63909E-07 |
| **Hippocampus** | | 7.28196E-09 | | 1.90775E-64 | | 5.81219E-10 | | 1.56169E-22 |
| **Amygdala** | | 0.005100329 | | 5.955E-57 | | 4.52848E-25 | | 6.13575E-13 |
| **Striatum** | | 0.018494101 | | 4.6602E-25 | | 7.73E-11 | | 6.18083E-05 |

**Number of Segment**

| **Area** | **Female control versa Male control** | **Female control versa Female PLX** | **Male control versa Male PLX** | **Female PLX versa Male PLX** |
| --- | --- | --- | --- | --- |
| **Cortex** | 0.181495104 | 8.32279E-32 | 2.79174E-17 | 1.15705E-12 |
| **Olfactory bulb** | 0.259432673 | 4.17813E-43 | 4.19607E-20 | 4.29791E-08 |
| **Cerebellum** | 2.4979E-08 | 0.000636165 | 1.8258E-06 | 8.95111E-05 |
| **Hippocampus** | 3.53358E-09 | 1.38403E-62 | 4.0334E-09 | 2.66004E-21 |
| **Amygdala** | 0.012198736 | 1.09764E-51 | 8.71499E-26 | 4.38264E-10 |
| **Striatum** | 0.02089021 | 7.84279E-25 | 1.65211E-10 | 4.50286E-05 |
| **Number Of Extremities** |  |  |  |  |
| **Area** | **Female control versa Male control** | **Female control versa Female PLX** | **Male control versa Male PLX** | **Female PLX versa Male PLX** |
| **Cortex** | 0.426569028 | 7.91686E-30 | 2.50976E-16 | 3.89412E-11 |
| **Olfactory bulb** | 0.245116391 | 1.40672E-37 | 1.53792E-19 | 2.18814E-05 |
| **Cerebellum** | 7.68918E-10 | 0.01772186 | 2.46424E-06 | 0.000586579 |
| **Hippocampus** | 3.46212E-08 | 9.97524E-59 | 1.09182E-09 | 2.7861E-18 |
| **Amygdala** | 0.014837858 | 2.19313E-46 | 6.16029E-22 | 3.09508E-09 |
| **Striatum** | 0.025884183 | 5.70877E-22 | 1.66788E-09 | 0.000474674 |

| **Density** |  |  |  |  |  |  |
| --- | --- | --- | --- | --- | --- | --- |
| **Males** | **Cortex** | **Hippocampus** | **Amygdala** | **striatum** | **Cerebellum** | **Olfactory bulb** |
| **Cortex** |  | 0.012311669 | 1.39128E-08 | 0.067174482 | 0.005118193 | 1.9732E-15 |
| **Hippocampus** | 0.012311669 |  | 2.00808E-05 | 0.114212007 | 7.23684E-06 | 1.5833E-13 |
| **Amygdala** | 1.39128E-08 | 2.00808E-05 |  | 2.37547E-07 | 3.87275E-11 | 1.84807E-08 |
| **striatum** | 0.067174482 | 0.114212007 | 2.37547E-07 |  | 5.89755E-06 | 1.3185E-13 |
| **Cerebellum** | 0.005118193 | 7.23684E-06 | 3.87275E-11 | 5.89755E-06 |  | 8.53606E-16 |
| **Olfactory bulb** | 1.9732E-15 | 1.5833E-13 | 1.84807E-08 | 1.3185E-13 | 8.53606E-16 |  |
| **Females** | **Cortex** | **Hippocampus** | **Amygdala** | **striatum** | **Cerebellum** | **Olfactory bulb** |
| **Cortex** |  | 0.000367897 | 0.065159985 | 5.76098E-08 | 1.25687E-09 | 1.52621E-10 |
| **Hippocampus** | 0.000367897 |  | 0.03061077 | 0.000810422 | 5.15008E-06 | 7.73132E-13 |
| **Amygdala** | 0.065159985 | 0.03061077 |  | 1.68424E-05 | 1.48991E-07 | 1.21065E-11 |
| **striatum** | 5.76098E-08 | 0.000810422 | 1.68424E-05 |  | 0.008490304 | 1.21802E-14 |
| **Cerebellum** | 1.25687E-09 | 5.15008E-06 | 1.48991E-07 | 0.008490304 |  | 3.42404E-16 |
| **Olfactory bulb** | 1.52621E-10 | 7.73132E-13 | 1.21065E-11 | 1.21802E-14 | 3.42404E-16 |  |
| **Total branches Length** |  |  |  |  |  |  |
| **Males** | **Cortex** | **Hippocampus** | **Amygdala** | **striatum** | **Cerebellum** | **Olfactory bulb** |
| **Cortex** |  | 0.00053949 | 0.113545664 | 2.29673E-07 | 2.02418E-29 | 1.0602E-15 |
| **Hippocampus** | 0.00053949 |  | 0.008002707 | 0.049743963 | 3.23741E-31 | 7.43908E-21 |
| **Amygdala** | 0.113545664 | 0.008002707 |  | 1.01135E-05 | 6.89236E-38 | 4.22664E-21 |
| **striatum** | 2.29673E-07 | 0.049743963 | 1.01135E-05 |  | 3.17421E-38 | 3.69385E-27 |
| **Cerebellum** | 2.02418E-29 | 3.23741E-31 | 6.89236E-38 | 3.17421E-38 |  | 0.000723871 |
| **Olfactory bulb** | 1.0602E-15 | 7.43908E-21 | 4.22664E-21 | 3.69385E-27 | 0.000723871 |  |
| **Females** | **Cortex** | **Hippocampus** | **Amygdala** | **striatum** | **Cerebellum** | **Olfactory bulb** |
| **Cortex** |  | 1.3232E-14 | 0.003613947 | 3.2344E-08 | 1.59067E-41 | 2.46128E-16 |
| **Hippocampus** | 1.3232E-14 |  | 6.44659E-09 | 0.033411594 | 8.3551E-65 | 9.53954E-44 |
| **Amygdala** | 0.003613947 | 6.44659E-09 |  | 0.000308527 | 6.59309E-65 | 1.11211E-30 |
| **striatum** | 3.2344E-08 | 0.033411594 | 0.000308527 |  | 4.59964E-47 | 5.48827E-30 |
| **Cerebellum** | 1.59067E-41 | 8.3551E-65 | 6.59309E-65 | 4.59964E-47 |  | 5.84721E-16 |
| **Olfactory bulb** | 2.46128E-16 | 9.53954E-44 | 1.11211E-30 | 5.48827E-30 | 5.84721E-16 |  |
| **Number Of Roots** |  |  |  |  |  |  |
| **Males** | **Cortex** | **Hippocampus** | **Amygdala** | **striatum** | **Cerebellum** | **Olfactory bulb** |
| **Cortex** |  | 0.002797316 | 0.000249575 | 0.001868477 | 2.51495E-18 | 2.48814E-06 |
| **Hippocampus** | 0.002797316 |  | 0.493718166 | 0.457777687 | 6.72197E-24 | 9.82722E-12 |
| **Amygdala** | 0.000249575 | 0.493718166 |  | 0.454611283 | 1.79418E-35 | 1.01936E-18 |
| **striatum** | 0.001868477 | 0.457777687 | 0.454611283 |  | 2.03902E-27 | 1.79101E-13 |
| **Cerebellum** | 2.51495E-18 | 6.72197E-24 | 1.79418E-35 | 2.03902E-27 |  | 3.84886E-07 |
| **Olfactory bulb** | 2.48814E-06 | 9.82722E-12 | 1.01936E-18 | 1.79101E-13 | 3.84886E-07 |  |
| **Females** | **Cortex** | **Hippocampus** | **Amygdala** | **striatum** | **Cerebellum** | **Olfactory bulb** |
| **Cortex** |  | 1.16842E-05 | 0.451693433 | 1.5795E-05 | 4.50645E-40 | 2.6707E-10 |
| **Hippocampus** | 1.16842E-05 |  | 4.33518E-06 | 0.415111777 | 1.48557E-53 | 9.71391E-23 |
| **Amygdala** | 0.451693433 | 4.33518E-06 |  | 7.43571E-06 | 8.41635E-49 | 1.18479E-12 |
| **striatum** | 1.5795E-05 | 0.415111777 | 7.43571E-06 |  | 1.05693E-45 | 2.88873E-20 |
| **Cerebellum** | 4.50645E-40 | 1.48557E-53 | 8.41635E-49 | 1.05693E-45 |  | 7.99237E-18 |
| **Olfactory bulb** | 2.6707E-10 | 9.71391E-23 | 1.18479E-12 | 2.88873E-20 | 7.99237E-18 |  |
| **Number Branch Points** |  |  |  |  |  |  |
| **Males** | **Cortex** | **Hippocampus** | **Amygdala** | **striatum** | **Cerebellum** | **Olfactory bulb** |
| **Cortex** |  | 9.12444E-05 | 0.091183052 | 1.04692E-05 | 1.96314E-31 | 1.3683E-09 |
| **Hippocampus** | 9.12444E-05 |  | 0.002854753 | 0.305315366 | 9.2749E-33 | 3.09348E-16 |
| **Amygdala** | 0.091183052 | 0.002854753 |  | 0.000476691 | 1.75823E-39 | 2.37045E-13 |
| **striatum** | 1.04692E-05 | 0.305315366 | 0.000476691 |  | 9.398E-35 | 9.99579E-18 |
| **Cerebellum** | 1.96314E-31 | 9.2749E-33 | 1.75823E-39 | 9.398E-35 |  | 5.93133E-08 |
| **Olfactory bulb** | 1.3683E-09 | 3.09348E-16 | 2.37045E-13 | 9.99579E-18 | 5.93133E-08 |  |
| **Females** | **Cortex** | **Hippocampus** | **Amygdala** | **striatum** | **Cerebellum** | **Olfactory bulb** |
| **Cortex** |  | 7.07412E-15 | 0.018850437 | 2.04186E-06 | 3.96009E-38 | 1.62504E-11 |
| **Hippocampus** | 7.07412E-15 |  | 2.80527E-11 | 0.002388912 | 5.94995E-64 | 6.07646E-40 |
| **Amygdala** | 0.018850437 | 2.80527E-11 |  | 0.000679231 | 1.66407E-66 | 8.94064E-24 |
| **striatum** | 2.04186E-06 | 0.002388912 | 0.000679231 |  | 7.3896E-44 | 9.44197E-24 |
| **Cerebellum** | 3.96009E-38 | 5.94995E-64 | 1.66407E-66 | 7.3896E-44 |  | 3.83283E-23 |
| **Olfactory bulb** | 1.62504E-11 | 6.07646E-40 | 8.94064E-24 | 9.44197E-24 | 3.83283E-23 |  |
| **Number Of Segments** |  |  |  |  |  |  |
| **Males** | **Cortex** | **Hippocampus** | **Amygdala** | **striatum** | **Cerebellum** | **Olfactory bulb** |
| **Cortex** |  | 2.98806E-05 | 0.056774202 | 9.27465E-06 | 2.88901E-32 | 7.4098E-08 |
| **Hippocampus** | 2.98806E-05 |  | 0.002185278 | 0.377794456 | 2.40175E-35 | 1.4683E-15 |
| **Amygdala** | 0.056774202 | 0.002185278 |  | 0.000821102 | 3.7054E-42 | 5.2596E-12 |
| **striatum** | 9.27465E-06 | 0.377794456 | 0.000821102 |  | 3.03084E-36 | 3.19565E-16 |
| **Cerebellum** | 2.88901E-32 | 2.40175E-35 | 3.7054E-42 | 3.03084E-36 |  | 2.45765E-10 |
| **Olfactory bulb** | 7.4098E-08 | 1.4683E-15 | 5.2596E-12 | 3.19565E-16 | 2.45765E-10 |  |
| **Females** | **Cortex** | **Hippocampus** | **Amygdala** | **striatum** | **Cerebellum** | **Olfactory bulb** |
| **Cortex** |  | 2.68645E-16 | 0.015563001 | 1.26903E-06 | 4.34358E-40 | 2.05102E-09 |
| **Hippocampus** | 2.68645E-16 |  | 1.93073E-12 | 0.000788858 | 6.72253E-66 | 2.24152E-38 |
| **Amygdala** | 0.015563001 | 1.93073E-12 |  | 0.000576543 | 1.79847E-69 | 2.70572E-20 |
| **striatum** | 1.26903E-06 | 0.000788858 | 0.000576543 |  | 4.97708E-45 | 1.08184E-21 |
| **Cerebellum** | 4.34358E-40 | 6.72253E-66 | 1.79847E-69 | 4.97708E-45 |  | 1.04573E-28 |
| **Olfactory bulb** | 2.05102E-09 | 2.24152E-38 | 2.70572E-20 | 1.08184E-21 | 1.04573E-28 |  |

| **Number Of Extremities** |  |  |  |  |  |  |
| --- | --- | --- | --- | --- | --- | --- |
| **Males** | **Cortex** | **Hippocampus** | **Amygdala** | **striatum** | **Cerebellum** | **Olfactory bulb** |
| **Cortex** |  | 0.000160761 | 0.227389948 | 0.000127268 | 3.18074E-30 | 9.33984E-11 |
| **Hippocampus** | 0.000160761 |  | 0.000915295 | 0.469146722 | 7.43509E-34 | 7.59506E-18 |
| **Amygdala** | 0.227389948 | 0.000915295 |  | 0.000733106 | 1.91109E-37 | 8.41574E-14 |
| **striatum** | 0.000127268 | 0.469146722 | 0.000733106 |  | 1.86032E-34 | 4.85935E-18 |
| **Cerebellum** | 3.18074E-30 | 7.43509E-34 | 1.91109E-37 | 1.86032E-34 |  | 2.9345E-07 |
| **Olfactory bulb** | 9.33984E-11 | 7.59506E-18 | 8.41574E-14 | 4.85935E-18 | 2.9345E-07 |  |
| **Females** | **Cortex** | **Hippocampus** | **Amygdala** | **striatum** | **Cerebellum** | **Olfactory bulb** |
| **Cortex** |  | 3.59151E-16 | 0.013392231 | 1.17286E-06 | 1.60478E-41 | 1.05024E-10 |
| **Hippocampus** | 3.59151E-16 |  | 5.02274E-12 | 0.001000574 | 2.3543E-65 | 8.10194E-39 |
| **Amygdala** | 0.013392231 | 5.02274E-12 |  | 0.000683687 | 1.55482E-68 | 9.75674E-22 |
| **striatum** | 1.17286E-06 | 0.001000574 | 0.000683687 |  | 3.93222E-45 | 1.5217E-22 |
| **Cerebellum** | 1.60478E-41 | 2.3543E-65 | 1.55482E-68 | 3.93222E-45 |  | 3.32942E-24 |
| **Olfactory bulb** | 1.05024E-10 | 8.10194E-39 | 9.75674E-22 | 1.5217E-22 | 3.32942E-24 |  |

P>0.01, P<0.01
